# Implicit visuomotor adaptation remains limited after several days of training

**DOI:** 10.1101/711598

**Authors:** Sarah A. Wilterson, Jordan A. Taylor

## Abstract

Learning in sensorimotor adaptation tasks has been viewed as an implicit learning phenomenon. The implicit process affords recalibration of existing motor skills so that the system can adjust to changes in the body or environment without relearning from scratch. However, recent findings suggest that the implicit process is heavily constrained, calling into question its utility in motor learning and the theoretical framework of sensorimotor adaptation paradigms. These inferences have been based mainly on results from single bouts of training, where explicit compensation strategies, such as explicitly re-aiming the intended movement direction, contribute a significant proportion of adaptive learning. It is possible, however, that the implicit process supersedes explicit compensation strategies over repeated practice sessions. We tested this by dissociating the contributions of explicit re-aiming strategies and the implicit process over five consecutive days of training. Despite a substantially longer duration of training, the implicit process still plateaued at a value far short of complete learning and, as has been observed in previous studies, was inappropriate for a mirror reversal task. Notably, we find significant between subject differences that call into question traditional interpretation of these group-level results.

**Significance Statement:** In this set of studies, we find that the implicit process cannot fully account for learning in adaptation tasks, such as the visuomotor rotation and mirror reversal tasks, even following several days of training. In fact, the implicit process can be counterproductive to learning. Most notably, we find significant between subject differences that call into question traditional interpretation of these group-level results.

## Introduction

A significant perturbation must be overcome when using a computer mouse to navigate around a monitor screen. If sliding the hand forward moves the cursor up the screen, a 90-degree rotation has been applied to the visual feedback of the hand. Despite this discrepancy, we do not feel as though we are explicitly compensating for the sift between hand and computer mouse. The movement is made automatically, with little conscious effort necessary to achieve the desired goal. Intuitively, this suggests an important role for the implicit process, compensation without strategic intervention, in motor skill. Recent research has focused on characterizing the mechanisms giving rise to the implicit processes used in motor skill learning, mostly through sensorimotor adaptation paradigms. A common approach is to ask participants to make point-to-point reaching movements to nearby targets while the hand is obscured from view and visual (or proprioceptive) feedback is artificially perturbed (Held and Schlank, 1959; Cunningham, 1989; Imamizu et al., 1995; Pine and Krakauer, 1996; Krakauer, 2009).

The results of these sensorimotor adaptation experiments have been interpreted under the framework that the implicit process is driving a significant proportion of learning (e.g. Scheidt, Dingwell, and Mussa-Ivaldi, 2001; Baddeley, Ingram, and Miall, 2003; Fine and Thoroughman 2006, 2007; Wei and Körding, 2009). However, a number of more recent studies have challenged this view. Initially, these studies disassociated explicit compensation strategies and the implicit process with post-experimental assays and questionnaires (Heuer and Hegele, 2008; Hegele and Heuer, 2010), instruction (Mazzoni and Krakauer, 2006; Benson et al, 2011), and modeling (Taylor and Ivry, 2014). This important early work showed that the slow, gradual learning of the implicit process is not necessary to compensate for visuomotor rotations.

In parallel to the recognition that strategies play an important role in adaptation tasks, we see rapidly emerging evidence that the implicit process is highly stereotyped regardless of the particular task demands. The implicit process asymptotes to a value far less than complete learning and is largely insensitive to perturbation magnitude (Bond and Taylor, 2015; Morehead et al., 2017). Morehead and colleagues (2017), revealed a stereotyped implicit learning curve when the incentive to strategize was completely removed. Surprisingly, asymptotic implicit learning remained insufficient to account for the perturbation after many hours of practice. (Morehead and Smith, 2017, Morehead et al., 2017; Kim et al., 2018). Finally, recent evident suggests that the contribution of the implicit process decreases with extended training (Avraham, et al., 2020).

The essential role of the implicit process may be in fine adjustments to very small errors. A few recent reports show that the implicit process is proportional for rotations under approximately 8 degrees and there may be an effect of reward (Kim, Morehead et al., 2018; Kim et al., 2019; Leow et al., 2018; Hutter and Taylor 2018). Given all of this, it is difficult to see the role of asymptotically constrained the implicit process in long-term motor learning.

However, a rest period, following exposure to a perturbed training environment, allows for continued learning in the absence of the stimuli (Brashers-Krug, Shadmehr, and Bizzi, 1996). If a rest period is important for learning, a single session may be insufficient to determine how the implicit process contributes over the long-term. The full effect of adaptation seen by Stratton (1896, 1897) and Kohler (1941, 1951a, 1951b), complete adaptation to an inverted world without need for explicit compensation, occurred following five full days of continuous exposure. Implicit learning may require extensive consolidation periods in order to fully compensate for a large visuomotor rotation (Krakauer and Shadmehr, 2006; Criscimagna-Hemminger and Shadmehr, 2008; but see Caithness et al. 2004).

In Experiment One, we investigate adaptative learning to a visual perturbation over five days of training. If the implicit process, as measured in this task, will eventually contribute substantially to overall compensation, we expect to see a steady increase in the proportion of learning accounted for by the implicit process over time. We find that the implicit process, on average, does not change over the training period. However, we also find substantial between-subject differences that raise questions about the validity of the measure.

An additional complication, not considered in Experiment One, is the directional nature of the implicit process. The implicit response to directional information appears to be automatic: the motor system adapts in a direction opposite to the perturbation even when such adaptation is task-irrelevant (Schaefer, Shelly, and Thoroughman, 2012, Morehead et al 2017; Butcher and Taylor, 2018). This response to directional information has led researchers to question the role that the implicit process might play in de-novo skill learning, such as the visuomotor mirror reversal task. This task requires a directional adaptation response *opposite* to that seen in rotation tasks (Figure 1; Gritsenko and Kalaska, 2010; Lillicrap et al., 2013; Telgen, Parvin, and Diedrichsen, 2014; Krakauer, 2009).

**Figure 1.**
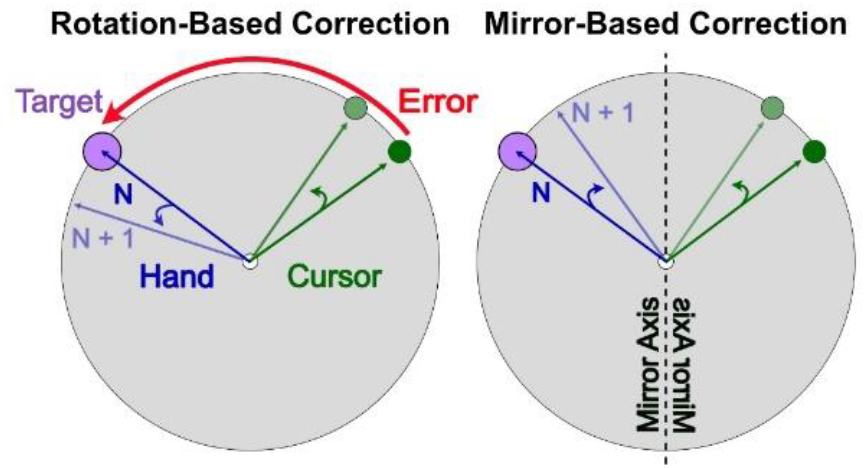
Appropriate correction for both rotation and mirror reversal perturbations. If the goal is to reduce error between the cursor and the target, the adaptive correction, denoted by the blue arrow, is counterclockwise for the rotation-based correction but clockwise for the mirror-based correction. Prior evidence supports the implicit process being appropriate for the rotation, but not for the mirror. This has been used as evidence that visuomotor rotation tasks produce adaptations of an established skill (reaching), while mirror-reversal tasks are a de-novo skill that do not benefit from adaptation.

Indeed, there is evidence that, even after a few days of training, the implicit process works against performance in mirror reversal tasks (Telgen et al, 2014; Kasuga et al, 2015). However, complicating this picture, a mirror reversal task is exactly what bilateral hippocampal lesion patient H.M. was able to complete – presumably without the benefit of explicit processes (Corkin, 1968). How was this accomplished if the implicit process – as we know it from previous visuomotor rotation studies – is unable to solve a mirror reversal?

In Experiment Two, we sought to determine if sufficient practice would transform the mirror-reversal task from one of *de novo* learning, to an adaptation task. That is, if the implicit process could become useful in a mirror task given a sufficient training period. This would give deeper insight into the role of the implicit process in early and long-term learning of novel motor skills. We found that, on average, the implicit process gradually decreases over time – slowly becoming more appropriate for the mirror reversal task. However, we again find substantial between-subject differences.

## Materials and Methods

In two experiments, twenty-four subjects participated in a visuomotor rotation task for one hour each day for five consecutive days. Subjects attempted to compensate for a visual perturbation of 45° while reaching to targets that appeared on a circle centered on the start-position.

### Participants

Seventeen first year graduate students, undergraduate students, and community members were recruited to participate in Experiment One. Of the seventeen, 3 subjects were removed from the experiment due to equipment failure and 2 additional subjects dropped-out of the experiment without completing all five days. The remaining twelve subjects [5 female, age 24.04 ± 2.11 yrs.] successfully completed the full experiment and their data is reported here. To have an equivalent number of subjects in the second experiment, fourteen subjects were recruited in the same manner as in Experiment One. Two of these subjects retired from the experiment before completion. Data is presented for the remaining twelve subjects [8 female, age 23.76 ± 3.04 yrs.].

Subjects were verified to be right handed by the Edinburgh Handedness Inventory (Oldfield, 1971) and self-reported having normal or corrected-to-normal vision. The experimental protocol was approved by the [Author University] Institutional Review Board and all subjects provided written, informed consent. All subjects received monetary compensation for their participation.

We did not have an *a priori* effect size estimate with which to calculate a power analysis. However, assuming a 3-degree change in mean and a SD of 3-degrees results in a suggested sample of 10 participants to achieve 90% power. A post-hoc power analysis conducted on the reduction of implicit adaptation found in Experiment 2 results in a power level of just over 85% to observe the reported effect.

### Apparatus

Subjects performed horizontal movements in a center-out reaching task similar to that first described in Bond and Taylor, 2017 (Figure 2). Stimuli were displayed on a 60 Hz, 17-in., Planar touch sensitive monitor (Planar Systems, Hillsboro, Oregon) and computed by a Dell OptiPlex 7040 machine (Dell, Round Rock, Texas) running Windows 7 (Microsoft Co., Redmond, Washington). Movements were recorded with a Wacom magnetic digitizing pen and tablet (Wacom Co., Kazo, Japan). Aiming locations were recorded by tapping the touch sensitive monitor, which was placed 25 cm above the Wacom tablet and obscured visual access to the right hand. The game was controlled by custom software coded in MATLAB (The MathWorks, Natick, Massachusetts), using Psychtoolbox extensions (Brainard, 1997, Kleiner, Brainard, and Pelli, 2007).

**Figure 2.**
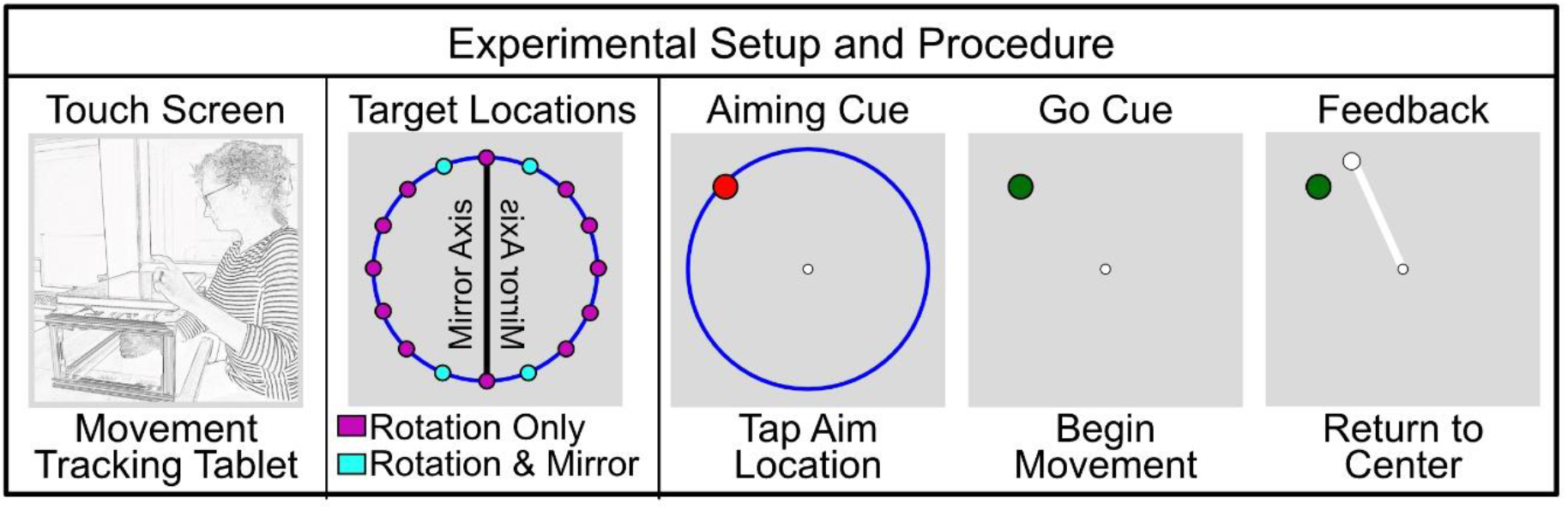
Experimental paradigm for assaying the implicit process to an imposed visual perturbation. The target locations for the mirror task were chosen to maintain a 45° perturbation (±22.5° from the mirror axis). Participants indicate their intended movement direction (re-aiming) by tapping on the “aiming ring”, which is shown here in blue. They then use the right hand to reach for the location that they think will bring the cursor to the target (ostensibly the re-aimed location). Any learning not accounted for by explicit re-aiming is presumed to be the implicit process.

### Procedure

In order to assay the operation of explicit strategies and the implicit process on a trial-by-trial basis, we asked participants to report their intended aiming location (i.e. the direction that they intend to move in order to compensate for the perturbation) by tapping the surface of a touch screen (following Bond and Taylor, 2017; Hutter and Taylor, 2018). Participants reported their intended aiming location and then make a reaching movement. The difference between the intended movement (explicit re-aiming) and the actual reach direction reveals any learning of which the subject was unaware (the implicit process).

Each trial began with subjects holding their right hand in the center of the workspace for 300 ms. Then, a circular orange target (0.25 cm radius) appeared 7 cm from the start position on a blue, 7 cm radius “aiming ring”. To begin a trial, the subject indicated their intended reach direction by tapping on the aiming ring with their left hand. Once an aim was recorded, the target turned from orange to green, the aiming ring disappeared, and the subject was able to begin their reach with the right hand (see Figure 2). Subjects were instructed to proceed through the experiment as quickly as possible, however, there was no time limit for tapping the aim location or beginning the reaching movement.

If a subject attempted to begin moving their right hand before an aim location was registered, the message “Remember to report aim” was displayed and the trial restarted. While reaching, subjects were instructed to move as quickly as possible past their intended aim location and back to the center in one smooth movement.

This “out-and-back” motion was encouraged in order to decrease the time necessary to return to the center of the workspace. After a successful movement beyond 7 cm, participants were guided back to the start position by a white ring. The ring was centered on the start position and its radius represented the distance of the subjects’ hand to the start position. However, subjects became highly accurate at stopping their out-and-back movement directly on the center of the workspace and used the guiding ring less overtime. This greatly increased the number of trials that could be completed in each one-hour session.

During each trial, once the right hand moved out past 7 cm, endpoint feedback was displayed for 500 ms in a position contiguous with the last point of on-line feedback. If the position of the cursor completely overlapped with the target (< 1 deg. of angular deviation), the subject heard a pleasant “ding”. Otherwise, an unpleasant “buzz” sounded. The feedback “Too Slow” was given if the reach time from start to 7 cm exceeded 600 ms (<1% of trials, these trials were excluded from further analysis).

A familiarization period of 16 trials with online feedback began the experiment. Two baseline periods followed the familiarization. The first baseline period was composed of 80 trials without feedback to determine if any subject held strong biomechanical biases that would not be averaged out by the large target set. The second baseline period provided veridical feedback, was also 80 trials long, and was designed to wash-out any drift that may have developed in the no-feedback phase. Subjects were not asked to report their aim during these baseline trials as it was assumed that they would always be aiming at the target.

A pause was included after the baseline trials so that the experimenter could explain the aiming procedure to the subjects. Subjects were explicitly instructed to indicate their intended reach direction, not the target or the cursor position, by tapping on the aiming ring with their left hand. Following these instructions, subjects completed 16 trials with veridical feedback to familiarize themselves with the touch screen and aiming procedure. The first day concluded with 608 perturbation trials, in which a 45° rotation between movements of the hand and cursor feedback was abruptly introduced. Subjects were provided with a three-minute break at the midpoint of training on each day.

Subjects were carefully observed during the familiarization trials and for the first 50 training trials to ensure that they understood the re-aiming instructions; further direction was provided as necessary during this time. If a subject was observed to ‘re-aim’ repeatably on the target, instead of adjusting their aim, the subject was stopped and the experimenter insured that they understood the purpose of the aiming procedure.

On days two through five, the perturbation was present from the first trial. Subjects were given a quick refresher of the aiming and movement instructions at the beginning of each day and were told that they would “pick up right where they left off” the day before. Subjects completed 800 trials each day, with an approximately 3-minute break halfway through. A total of 3,808 training trials were completed. At the end of day five, a 32-trial washout period was completed without feedback. During this washout, subjects were instructed to discontinue any strategy they had developed and aim straight for the target.In Experiment One (Rotation-Task), the perturbation was a 45° rotation in either the clockwise or counterclockwise direction (counterbalanced between subjects). In Experiment Two, the perturbation was a mirror about the vertical midline. In order to allow comparison between the two experiments, the four targets in Experiment Two (Mirror-Task) were located 22.5° from the midline so that the solution to the perturbed reach was a 45° angle. The 16 targets in Experiment One were evenly spaced 22.5° apart on the aiming ring (Figure 2). Both experiments were programmed such that each of the targets appeared before any was again repeated.

### Data and statistical analyses

The experiment presentation, data collection, and statistical analysis were all completed in Matlab (The Mathworks, 2016b). During both experiments, the digitizing tablet logged the trajectory of the right hand and the touchscreen monitor recorded the position tapped to indicate re-aiming location. To allow averaging across targets, hand trajectories were transformed to a common axis with the target at zero degrees. Additionally, the hand trajectories were transformed into heading angles by examining the average angle of the hand from a straight-line path to the target between 1 and 3 cm into movement. This procedure prevents the influence of visual feedback control on estimates of learning. An aiming angle for each trial was defined by the angle between the target and the location tapped on the touchscreen. The implicit process was calculated as the subtraction of the aim angle from the hand heading angle in each trial (Taylor, Krakauer, and Ivry, 2014). In all measures, a positive angle represents a counterclockwise divergence from the target. As we measured only two values, and computed the implicit process from those values, we conducted statistical analyses for only the aiming angles and the implicit process angles. We report the mean and standard deviation of hand angles for completeness only. Reaction times were calculated as the interval from target appearance to aim-report, except where noted. All data used in parametric statistical tests were tested for normality using Lilliefors test.

### Modeling

To predict the time courses of explicit and implicit processes in the mirror task, we used a modified version of the two-state model (Smith et al., 2006). Here, we modeled explicit re-aiming as the fast learning process (Xf) and the implicit process as the slow learning process (Xs) over the course of 200 trials (for a detailed account of modeling explicit/implicit learning as fast/slow learning, see previous work by McDougle, Bond, and Taylor 2015). In addition, we assumed that explicit re-aiming is updated based on target error, while the implicit process is updated based on the aim-to-cursor distance (Equations 1 and 2).

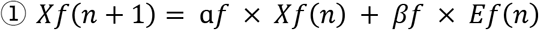

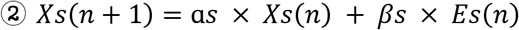

Where *Ef* is the error between the target and the cursor locations, while *Es* is the error between explicit re-aiming *(Xf)(n)* and the cursor location. Note, we did not input actual values of explicit re-aiming in these simulations but instead simply treated them as a faster learning process. In addition, we did not fully collapse the target locations to a common axis in order to demonstrate how the mirror-reversal perturbation causes different signed errors depending on target location. As such, we separately simulated target locations in the first and third quadrants, and second and fourth quadrants. The values on *αf, βf*, as, and *βs* were determined by hand tuning these parameters to the simple rotation case (Rotation-Task-Rotation-Correction Model, Figures 7B and 8B; *af* = 1, *βf* = 0.2, as = 0.98, and *βs* = 0.01). Note, we chose a value for as that was slightly lower than one to simulate an asymptotic value of the implicit process that is similar to what has been observed experimentally. These same parameter values were then used for both the Mirror-Task-Rotation-Correction and the Mirror-Task-Mirror-Correction Model. In the Mirror-Task-Rotation-Correction Model, the error for the implicit process was defined in exactly the same way as in the Rotation-Task-Rotation-Correction Model.

To demonstrate the effect of the implicit process switching the error-correction calculation to be consistent with the mirror reversal, we flipped the sign of the implicit error term in the Mirror-Task-Mirror-Correction Model (Equation 3).

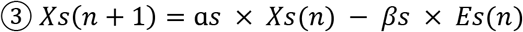

The modeling code described here is freely available online at [URL redacted for double-blind review]. The code is also available as Extended Data.

## Results

### Experiment One

In Experiment One, twelve subjects participated in a visuomotor rotation task for one hour each day for five consecutive days. Subjects attempted to compensate for a visual perturbation of 45° while reaching to targets that appeared on a circle centered on the start-position. To dissociate explicit and implicit learning, subjects reported their intended end position (re-aiming) using a touch-screen monitor (Bond and Taylor, 2017; Hutter and Taylor, 2018, Figure 2).

As expected, participants were able to compensate for the visual perturbation during the first day of training (Figure 3). Nearly perfect performance, as measured by the hand heading angle, was achieved by the end of the first day (44.8 ± 2.0°) and was maintained in the fifth day of training (44.7 ± 0.8°). Interestingly, this high level of performance was supported by a relative mixing of explicit re-aiming and the implicit process both on the first day (28.1 ± 6.7° re-aiming vs. 16.7 ± 6.4° adaptation) and on day five of the experiment (30.7 ± 16.2° re-aiming vs. 13.9 ± 16.3° adaptation). Note, while it appears that explicit re-aiming accounts for nearly twice as much learning as the implicit process, performing inferential statistics between explicit and implicit learning is theoretically inappropriate since our implicit measure is derived from our explicit measure.

**Figure 3.**
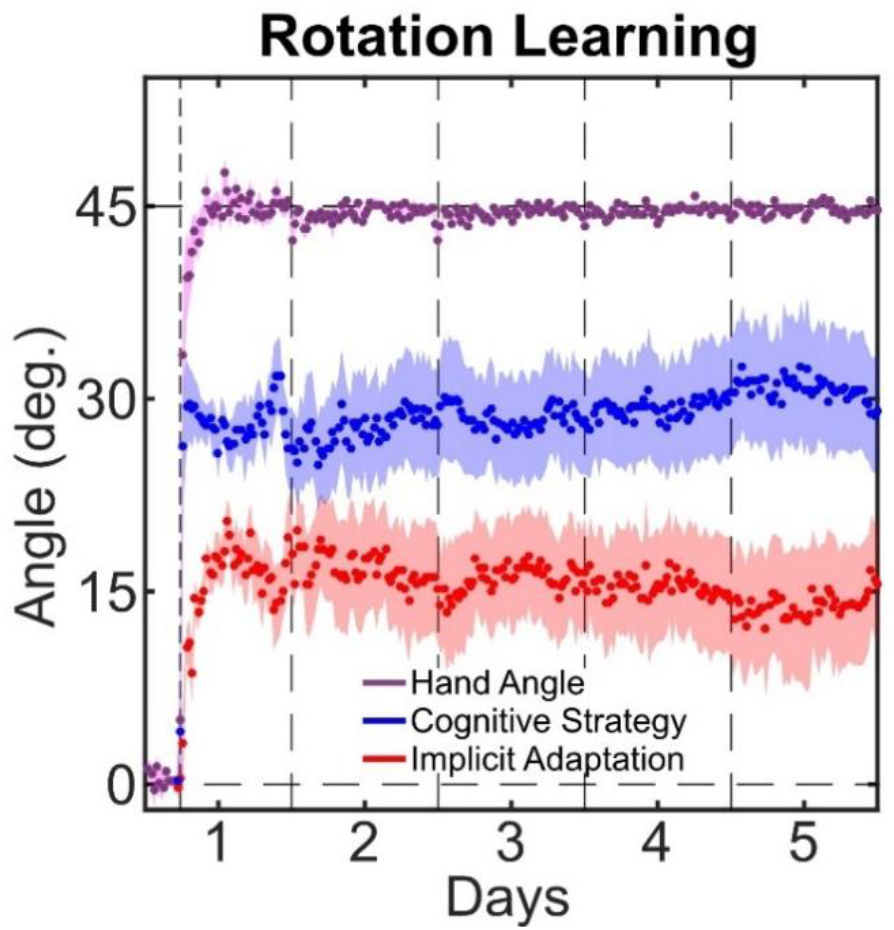
Time course of responses over five days of training to a 45° rotation. All responses were rotated to a common axis with the target at 0° and the solution at 45°. The vertical dashed lines represent first the onset of the rotational perturbation, then day breaks. Shaded areas represent between-subject standard error.

#### Analysis of within-subject and between-subject variance

While learning appears to be divided between explicit and implicit processes, visual inspection shows a notable increase in the between-subject variance for explicit re-aiming and the implicit process following the first day of training. This suggests that the relative contribution of explicit and the implicit process may differ dramatically between subjects. Indeed, if we sort the subjects by amount of the implicit process on Day 5 we find a full continuum of learning patterns (Figure 4). Our twelve subjects run the gamut from no the implicit process (Figure 4B), through a mixture of the implicit process and explicit re-aiming (Figure 4C), to complete the implicit process (Figure 4D). These results are consistent with the range of the implicit process previously found in numerous studies, including those using standard visuomotor rotation tasks without re-aiming reports (Taylor and Ivry, 2014), perturbations introduced in a pseudorandom walk (Stark-Inbar, 2017), and task-irrelevant-visual-error clamp (Morehead et al 2017; Kim et al 2018; Kim et al 2019).

**Figure 4.**
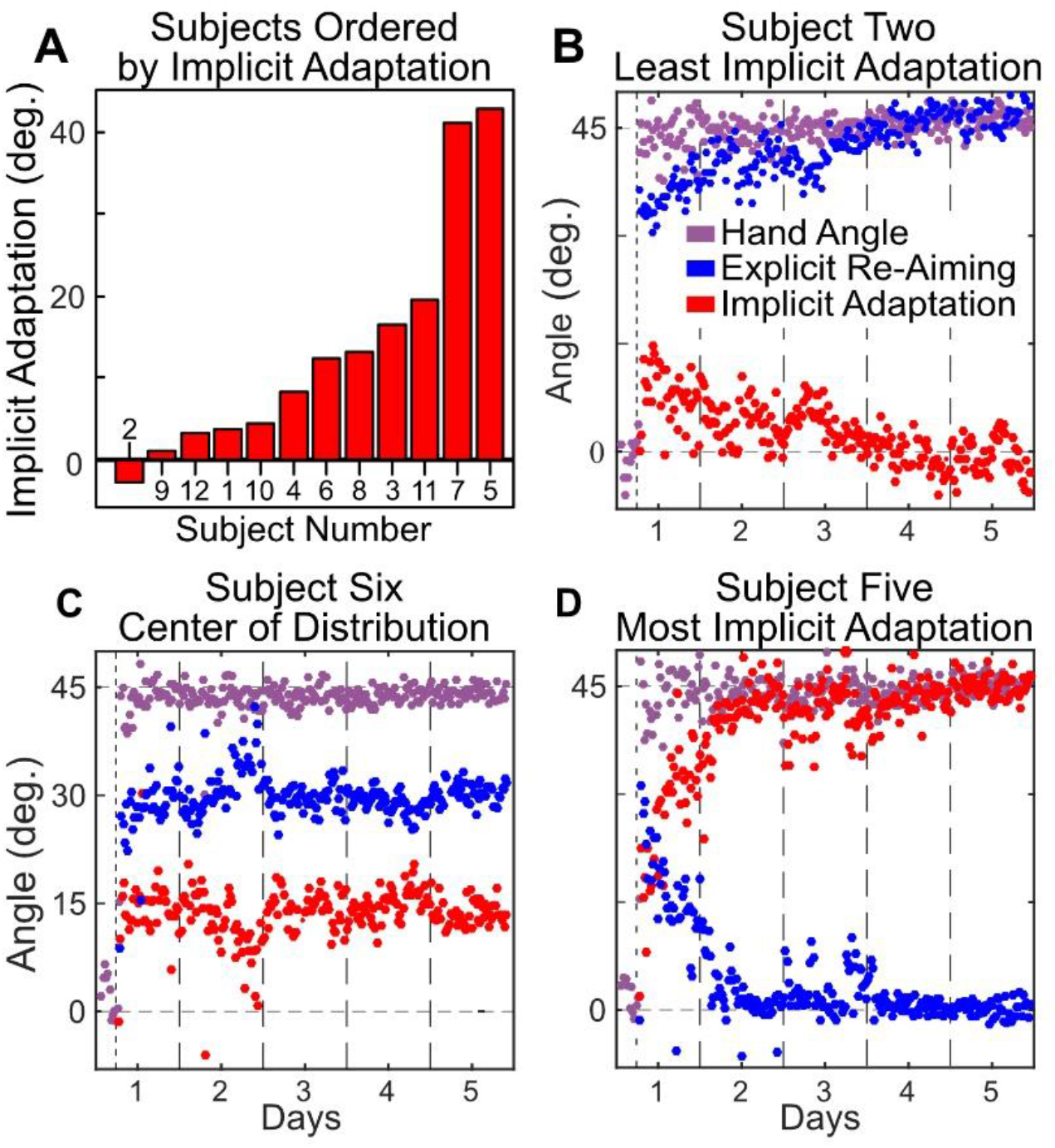
(A) A bar graph ordering subjects by the average amount of the implicit process as calculated from the last 200 trials of day five. (B) The subject (#2) reporting the most explicit re-aiming at the end of learning, assumed to have the least the implicit process (i.e. aiming directly at the solution with no perceptible the implicit process at the end of day five). (C) A subject (#6) taken from the center of the distribution. (D) The subject (#5) reporting the least explicit re-aiming (i.e. aiming at or very near to the target with all learning accounted for by the implicit process).

Despite these large between-subject differences, participants appeared to be remarkably consistent in maintaining their personal relative contribution of explicit and implicit learning within and across days. To quantify this, we performed a regression comparing explicit re-aiming and the implicit process at the end of each day to the same measure at the beginning of the following day (Figure 5). We found a high correlation between the end of one day and the beginning of the next for both explicit re-aiming (*Pearson’s r* = 0.94) and the implicit process (*r* = 0.94). To underscore that this correlation only reflects consistency at the individual level and not at the group level, we sought to quantify group consistency by shuffling the individual data. Here, we randomly assigned the prior days of each subject to the following days of a different subject 10,000 times. Averaging the correlation over these runs results in a correlation coefficient a fraction of the size seen for the true data (explicit re-aiming *average r* = −0.07; the implicit process *average r* = −0.07).

**Figure 5.**
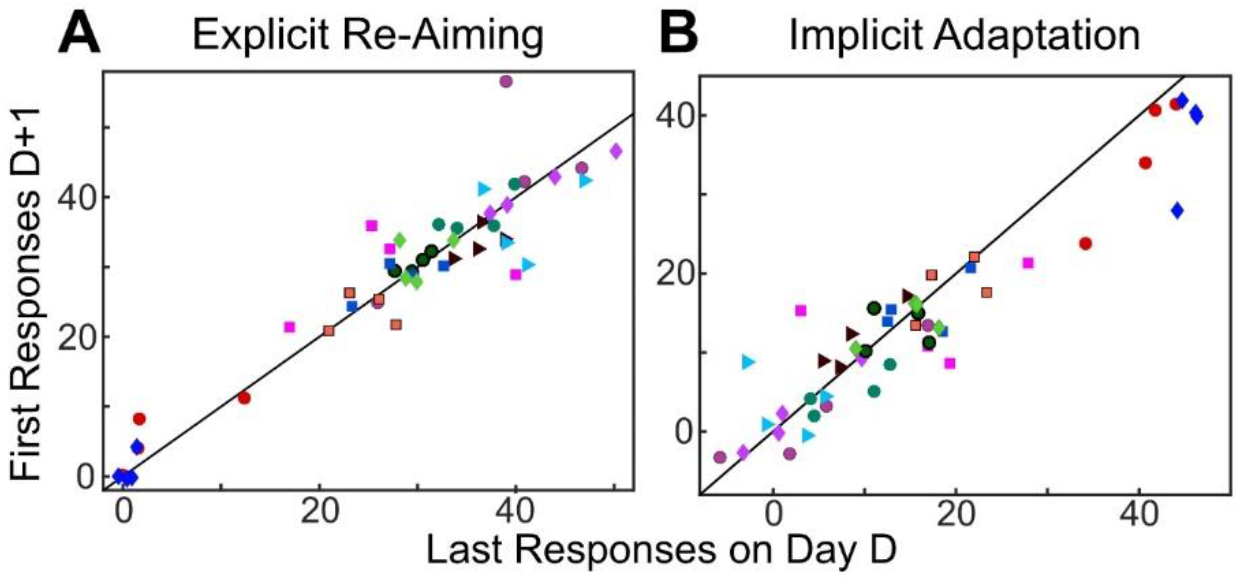
Average explicit re-aiming (A) and the implicit process (B) in last bin (16 trials) of day D plotted against first bin of day D+1 (*r* = 0.94). Each subject contributed 4 data points, and is represented by shape and color for visual clarity. The diagonal lines represent the unity line between responses on D and D+1. The marker color and shape represent an individual subject across all figures in this manuscript.

#### Analysis of adaptation measurements

When we use trial-by-trial aiming data to infer the implicit process levels, two subjects appear to compensate fully with the implicit process and five subjects appear to be using a mostly explicit strategy, with the five remaining subjects somewhere in-between. We compared this result with the traditional aftereffects measure, which was completed at the end of the 5^th^ day of training. During washout trials where subjects were instructed to aim directly at the target without using any strategy and on which no feedback was given, not a single subject shows a full 45° aftereffect, or a 0° aftereffect – which would have been indicative of learning entirely via the implicit process and explicit re-aiming strategies respectively (Figure 6A).

**Figure 6.**
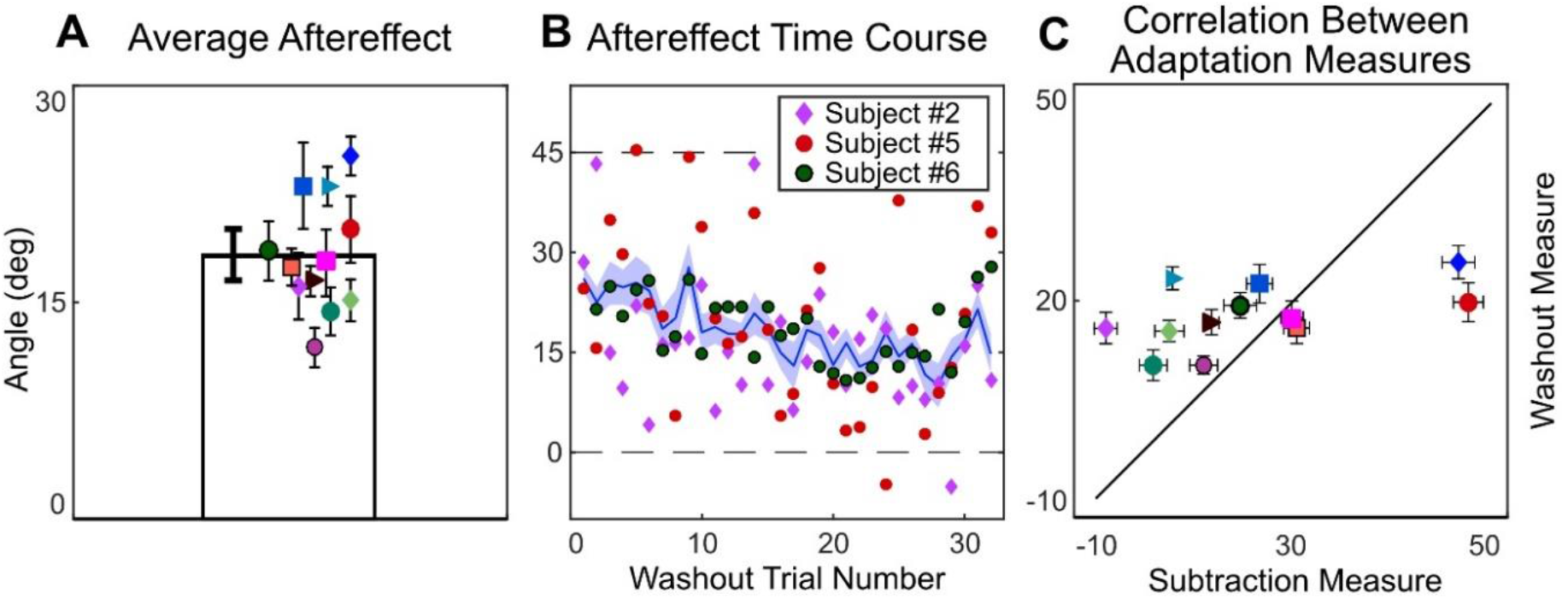
(A) Average aftereffect for each subject averaged across the 32 trials of no-feedback washout (with subjects instructed to aim and move directly to the target). Error bars represent one standard error of the mean. (B) The time course of washout for the average across subjects, and individual trial values for the three example subjects defined in Figure 3. Shaded area represents between-subjects standard error. (C) Correlation between each individuals’ adaptation as measured via subtraction (hand angle minus explicit re-aiming over the last 32 trials of training) vs washout (average hand angle in 32 washout trials). Error bars represent one standard error of the mean along each dimension. The marker color and shape represent an individual subject across all figures in this manuscript.

It is visually apparent that the aftereffects measure does not display the same range of adaptation that we saw using the aim subtraction method. To quantify this visual observation, we ran a correlation analysis comparing the implicit process in the final 36 trials of training (calculated with the subtraction measure) with the implicit process measured in the first 36 trials of the washout block (calculated by discrepancy from target location, with instruction to reach directly to target). We note that any correlation analyses on a limited sample size (N=12) should be viewed with caution. Additionally, it is important to note that we observed extremely variable trial-to-trial measurements within a subject. While the average adaptation, as reported in most studies of the implicit process, corresponds well to the mean of trial-by-trial adaptation calculated through the aiming subtraction method, individual measures do not (Figure 6B).

Confidence is warranted in suggesting that some participants fully strategized their way out of the rotation problem. This is because they had to be aware of the correct solution in order to tell us that they intended to move 45° away from the target. Contrast this with the fully implicit subject, who is telling us that they don’t perceive a rotation and so taps on or very near the target each trial. Such behavior could also be explained by a misinterpretation of the instructions or simple disregard for the instruction. Note, however, participants were required to report their intended aiming strategy on each trial, limiting the effort saved by reporting inaccurately. Additionally, if participants didn’t understand the instructions, we would expect that they never re-aimed or fully re-aimed, and that this behavior would be consistent from the introduction of the rotation. Instead, we see gradual changes with training. Participants who displayed full the implicit process followed a consistent time course where the implicit process gradually increased with training while their explicit re-aiming gradually decreased. Likewise, participants who displayed nearly full explicit re-aiming increased their re-aiming angle with training while the implicit process gradually decreased. Importantly, in post-participation debriefing one of the participants who displayed full the implicit process indicated that they thought the perturbation was removed; the second such participant declined to be interviewed.

We conducted several post-hoc analyses in an attempt to determine the cause of the variability in the implicit process found in this experiment. Regression analyses with day-five the implicit process regressed by inter-trial interval, reaction time, movement time, or reach end-point variability, all produced null results (*p* > .26). It is likely, however, that this is the result of limited power driven by a small sample size for this type of analysis. A study specifically designed – with much greater power – will be required to identify the source of individual variability.

#### Experiment One Conclusion

The average response to long-term adaptation in a visuomotor rotation tasks suggests that both the implicit process and explicit re-aiming continue to play a role in performance even after several days of training. On average, we found the implicit process to be constrained, which is consistent with the findings from studies examining the capacity of single-session implicit learning. However, we saw great variability between subjects, including evidence from two subjects that implicit recalibration could eventually replace explicit re-aiming.

We next investigate the usefulness of implicit recalibration processes in a *de novo* skill learning task, the mirror reversal task.

### Experiment Two

In Experiment Two, twelve subjects participated in a visuomotor mirror reversal task for five consecutive days. Subjects compensated for a mirror perturbation originating at the midline while reaching to targets that appeared on a circle around the start position. Targets were positioned such that the correct response resulted in a 45° angle between hand and feedback. Subjects reported intended re-aiming using a touch-screen monitor before each trial. The procedure was exactly the same as previously used in Experiment One.

As shown in Figure 1, useful recalibration in rotation paradigms has been described as that responding to the discrepancy between the cursor feedback and aiming location (Taylor and Ivry, 2011; Day, Roemmich, Taylor, and Bastian, 2016). For the implicit process to be useful in the mirror reversal task, the implicit error signal must be calculated in the opposite direction as it is for rotational perturbations (Figure 7). Additionally, the mirror reversal imposes a counterclockwise error in quadrants one and three but a clockwise error in quadrants two and four – similar to a dual adaptation paradigm (Howard et al., 2012; Schween et al., 2019). This complicates the transformation space over targets and contributes to the learning difficulty.

**Figure 7.**
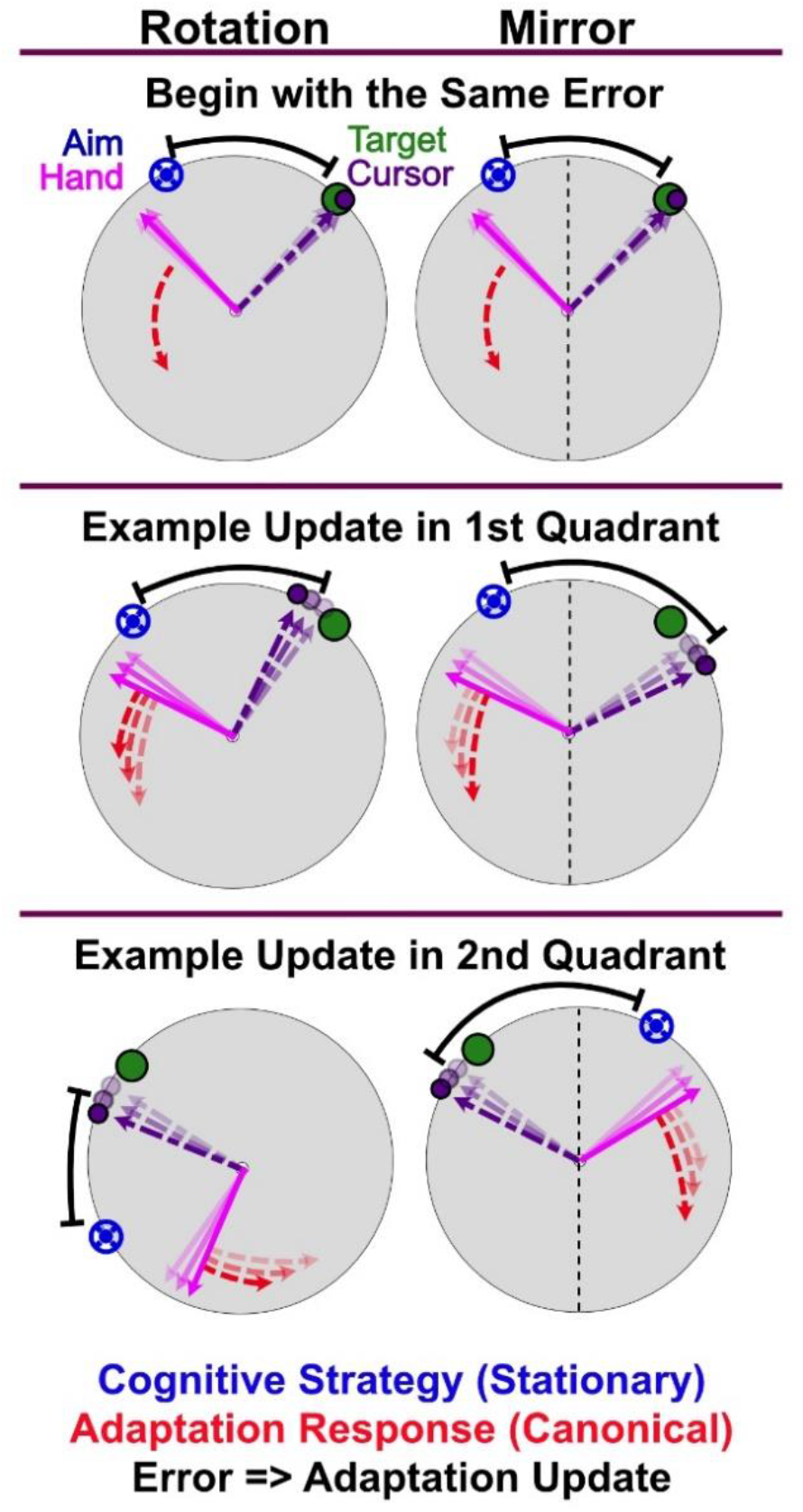
Six scenarios with hand angle (magenta) responding only to implicit recalibration (red). Explicit strategy (blue) is held constant to simplify the figure. The implicit process (red) is driven by the error (black) between the re-aimed location (blue) and feedback (purple). Additive adaptation pulls the hand away from the re-aimed location and results in decreased error for a rotational perturbation, and increased error under a mirror reversal. The error sign is flipped in quadrants two and four under a mirror reversal, but not a rotation. Greater transparency of the vectors and endpoints indicates trials further into the past.

For clarity of visualization on graphics, and to underscore the difficulty of accumulating the implicit process during a mirror task, we combined responses for the targets in quadrants 1 and 3 (i.e., at 67.5° with those at 247.5°) and responses for the targets in quadrants 2 and 4 (i.e., at 112.5° with those at 292.5°; Figure 7). Unlike in the rotation task, normalizing all target locations to zero does not assist in visualizing the data. Statistical analyses were conducted on the full data set, combining all target locations by flipping the sign of quadrants two and four prior to averaging.

Unlike the visuomotor rotation in Experiment One, participants on average slightly over-compensated for the mirror reversal perturbation on the first day (48.5 ± 3.8°), however, performance on the final day was nearly perfect (45.7 ± 1.3°, Figures 8A and 9A).

**Figure 8.**
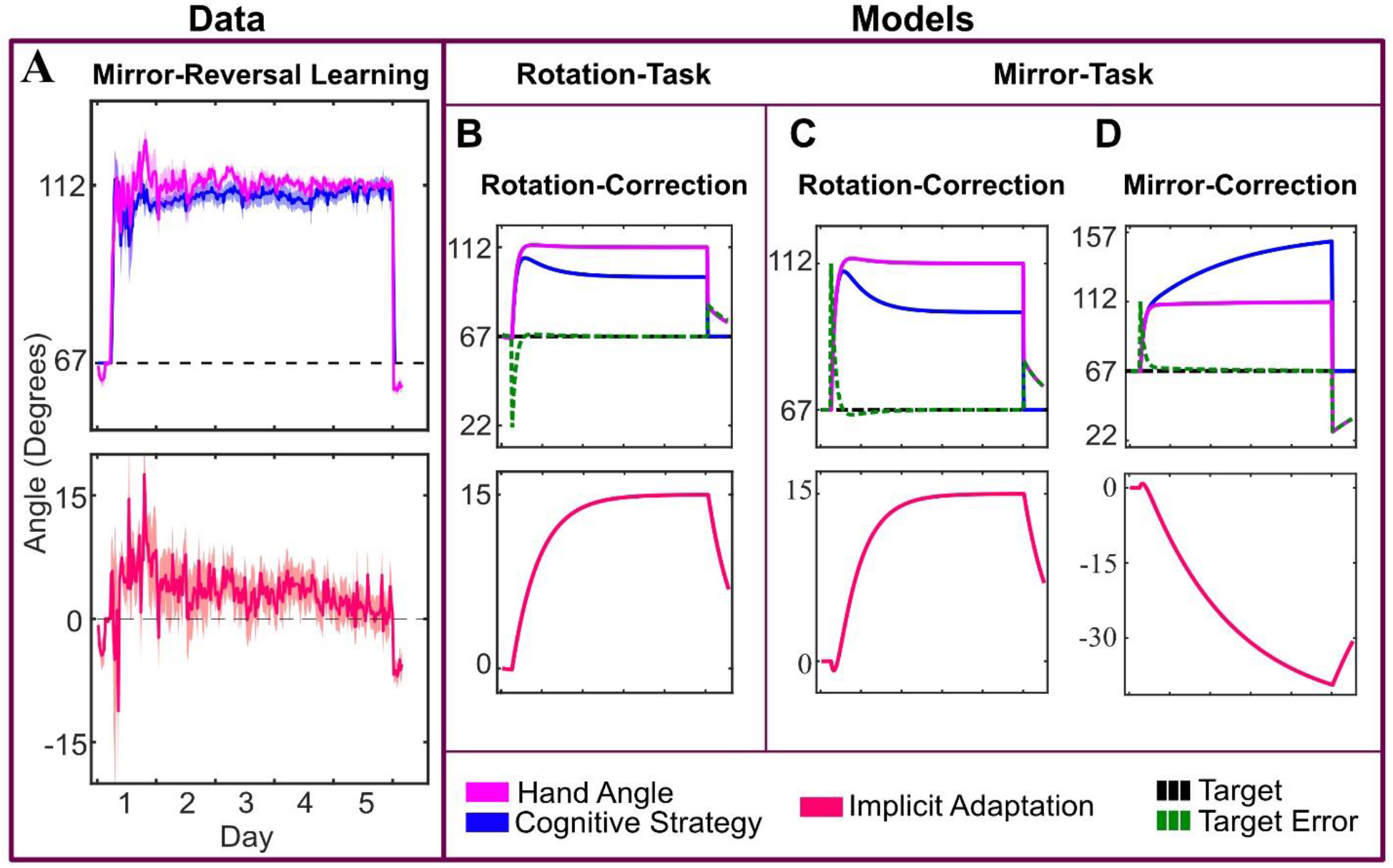
(A) Mirror reversal learning for all 12 subjects to targets in the first and third quadrants, normalized to 67.5°. The solution is 112.5°. Implicit recalibration is shown separately with positive values being appropriate for a rotational perturbation. (B) Rotation-Task-Rotation-Correction Model for target at 67.5° with parameters tuned to limit the implicit process to about 15°. (C) Mirror-Task-Rotation-Correction Model with parameters defined as in ‘B’. (D) Mirror-Task-Mirror-Correction Model with parameters defined as in previous. Implicit error term is reversed from the rotation correction.

**Figure 9.**
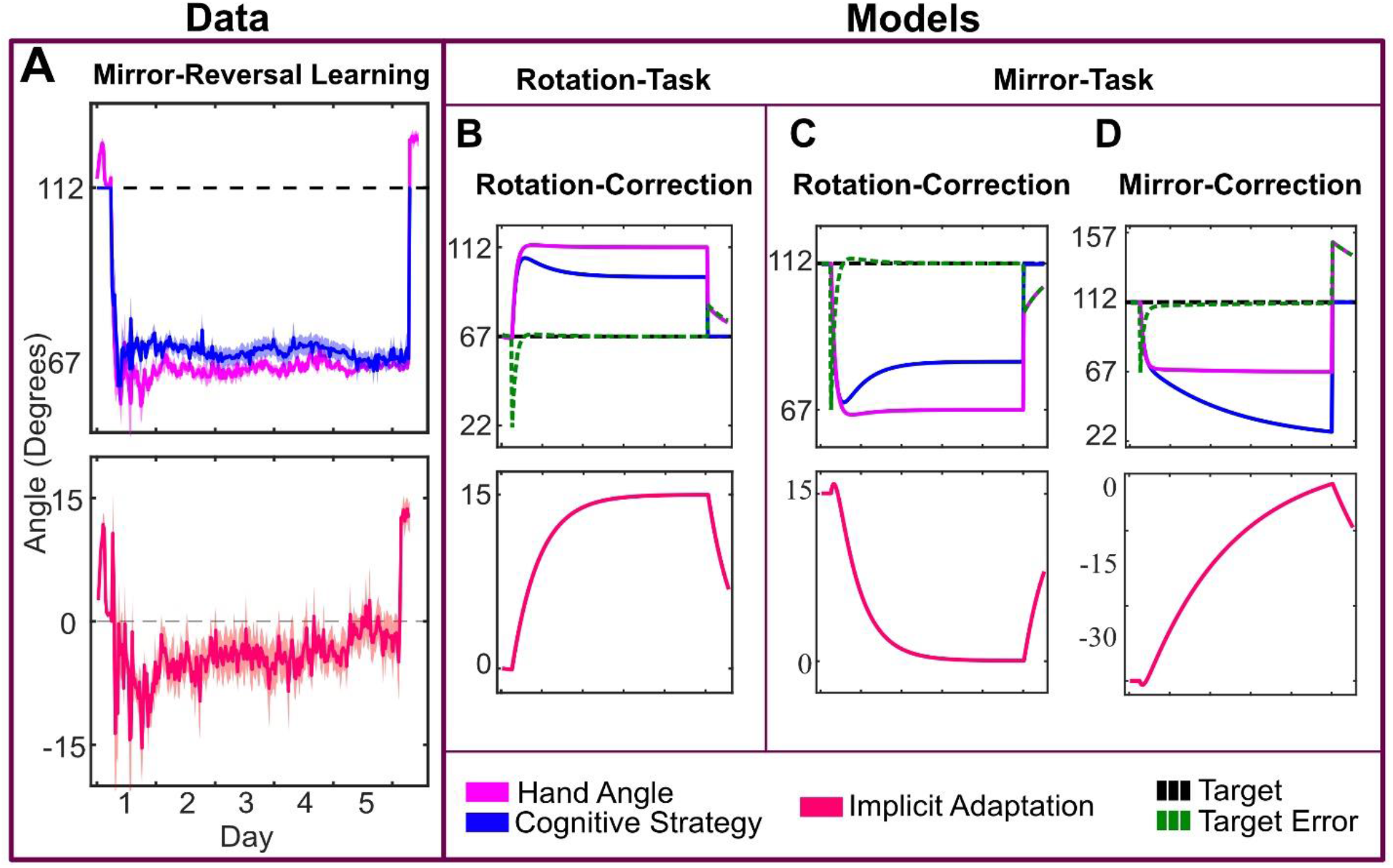
(A) Mirror reversal learning for all 12 subjects to targets in the second and fourth quadrants, normalized to 112.5°. The solution is 67.5°. Implicit recalibration is shown separately with negative values being consistent with a rotational perturbation. (B) Rotation-Task-Rotation-Correction Model for target at 112.5° with parameters tuned to limit the implicit process to about 15°. (C) Mirror-Task-Rotation-Correction Model with parameters defined as in ‘B’. (D) Mirror-Task-Mirror-Correction Model with parameters defined as in previous. Implicit error term is reversed from the rotation correction.

On the first day, performance was again a mixture of the implicit process (8.0 ± 4.6°) and explicit re-aiming (40.5 ± 5.4°). However, while the implicit process initially contributed to learning (t-test of first day, t = 6.050, p < .001), it gradually decreased over time (paired t-test between adaptation on first and last day, t = 2.202, p = .049) and was not significantly different from zero on the final day (2.2 ± 7.8°, t = 0.961, p = .357). On average, participants fully compensated for the mirror reversal with explicit re-aiming (43.6 ± 7.6°) on the final day of training. Assuming a constant rate of change and that the implicit process is capped at approximately 15° (as seen in Experiment One), the implicit process would become more appropriate (i.e., switch signs) for a mirror-reversal after 10-14 days.

#### Analysis of within-subject and between-subject variance

As with the visuomotor rotation, there was significant variance in how subjects responded to the visuomotor mirror reversal (approximately −15° to 15°, Figure 10). Subjects were successful through a combination of the implicit process and explicit re-aiming, lost all the implicit process by the end of training, or reversed the direction of the implicit process. Reversed, visualized as negative, implicit adaptation is appropriate for the mirror reversal task.

**Figure 10:**
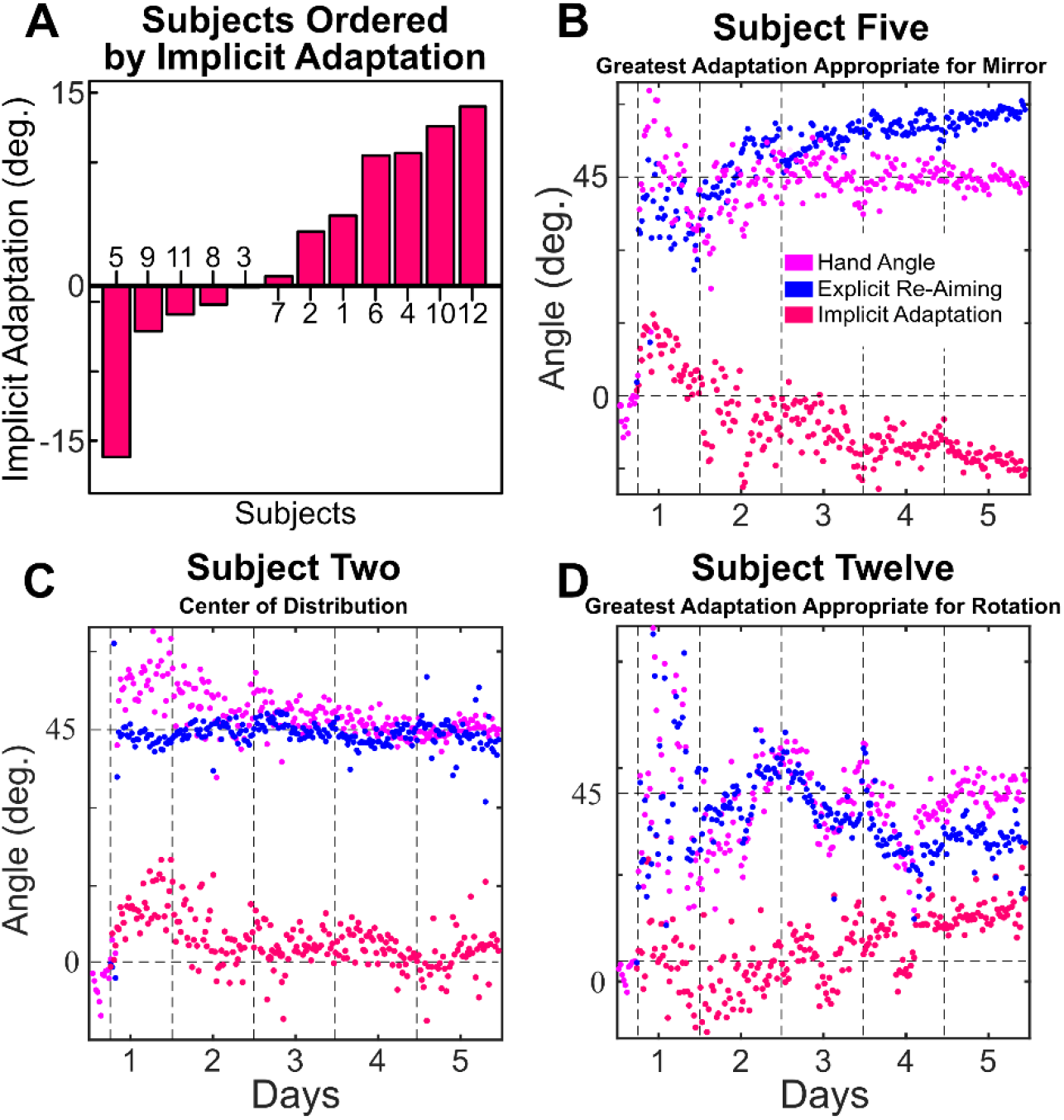
(A) A bar graph ordering subjects by the average amount of explicit re-aiming that they reported during the last 200 trials of day five. (B) The subject (#5) reporting to have reversed their implicit process so that it is appropriate for the mirror reversal at the end of day five. (C) A subject (#2) representative of the average taken from the middle of the distribution. (D) The subject (#12) reporting the most adaptation in the direction that is more appropriate for a rotation.

As in Experiment One, we conducted a linear regression analysis to determine how consistent subjects were across days (Figure 11). We again found remarkable consistency within each subject; expressed as a high correlation between the end of one day and the beginning of the next for both explicit re-aiming (*r* = 0.865) and the implicit process (*r* = 0.770). When this result was contrasted with a randomization between days D and D+1, the result is a far worse average correlation than the true data for both explicit re-aiming (*average r* = −0.067) and the implicit process (*average r* = −0.071). Again, while we note that correlations on small sample sizes should be viewed with caution, due to the extremely high correlational relationships we don’t think there is reason to believe that the inclusion of more participants would change the conclusion in this case.

**Figure 11.**
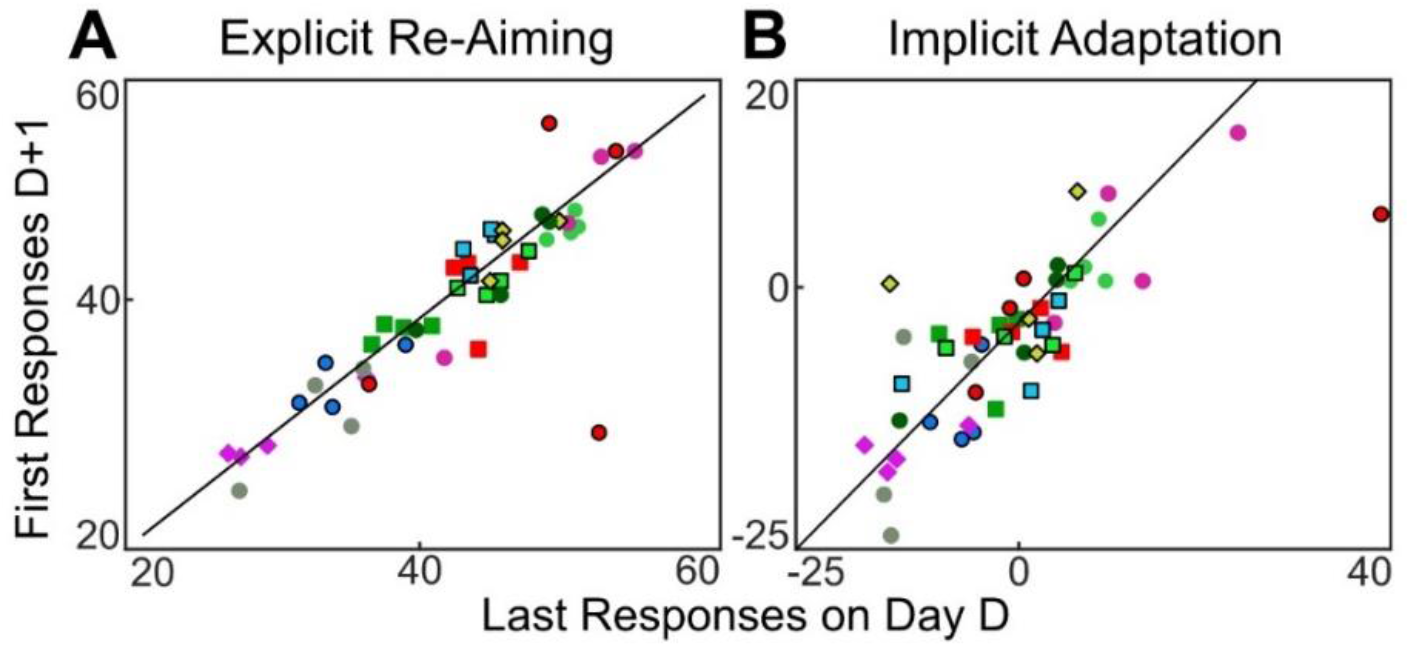
Average explicit re-aiming (A) and the implicit process (B) in last bin (16 trials) of day D plotted against first bin of day D+1 (*r* = 0.77 and r = 0.87, respectively). Each subject contributed 4 data points, and is represented by shape and color for visual clarity. The diagonal lines represent the unity line between responses on D and D+1.

**Figure 12.**
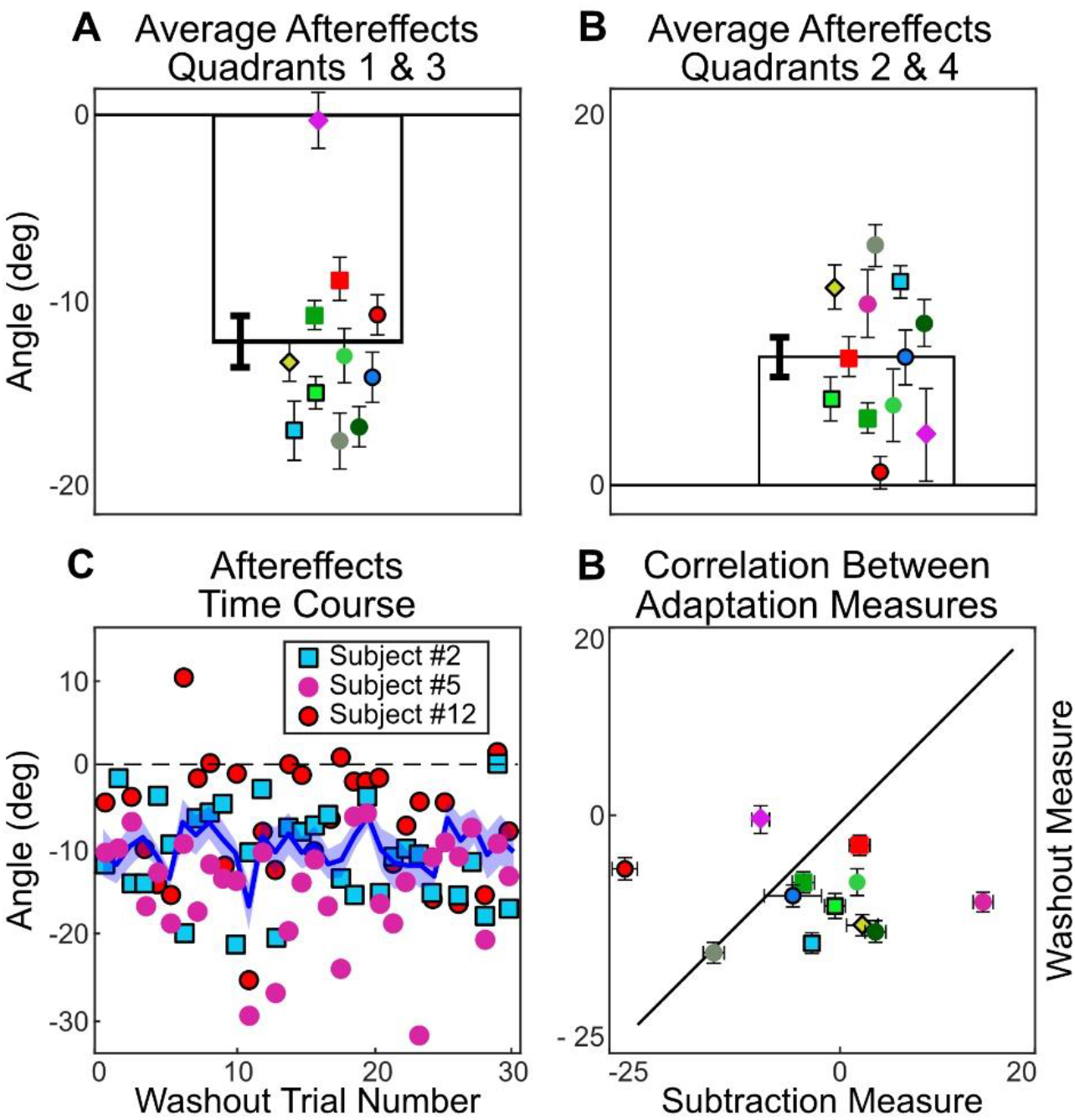
(A) Average corrected aftereffects for each subject across the 32 trials of no-feedback, washout (with subjects instructed to aim and move directly to the target). Each subject is jittered left to right in order of least to most explicit re-aiming at the end of Day 5. (B) The time course of washout for the average across subjects and individual trial values for three representative subjects [#5 (Lowest), #8 (Median), #12 (Highest), re-aiming on last day). The shaded area represents the standard error of the mean for all subjects.

#### Analysis of adaptation measurements

Mirror learning has been viewed as a skill learning process, as opposed to adaptation, because it does not result in the same response properties as a visuomotor rotation, such as rate of learning, offline consolidation, and shifts in a speed-accuracy tradeoff (Telgen, Parvin, and Diedrichsen, 2014). Given the complexity of forming an intuition about the implicit process in the mirror (Figure 7), we simulated the potential results using a modified two-state-space model (Smith et al., 2006). The goal was to form graphical representations of responses based on mirror-appropriate adaptation or adaptation to an alternating rotation. The two-state model characterizes learning in visuomotor rotation tasks in terms of two separate learning processes: one that learns slowly but retains memory and the other learns fast but quickly forgets. Recent work suggests that the slow process might be the implicit process while the fast process is explicit re-aiming (McDougle, Bond, and Taylor, 2015). We adopted the conventions of this model to predict learning based on the sign of the error signal. Note, our goal here is not to model mirror reversal learning *per se*, but to determine if the pattern of the implicit process we observed is more consistent with learning appropriate for a mirror reversal or for a visuomotor rotation.

Within the model, we assumed that the explicit re-aiming is updated based on target error, while the implicit process is updated based on the angle between the aim and the cursor (See methods for details). The learning and forgetting rates were determined by tuning these parameters to the simple rotation case in order to cap the implicit process at approximately 15° (Rotation-Task-Rotation-Correction Model, Figures 8B and 9B; Morehead and Smith, 2017). These same parameter values were then used for both of the Mirror-Models. In the Mirror-Task-Rotation-Correction Model, the error for the implicit process was defined in exactly the same way as in the Rotation-Task-Rotation-Correction Model (Figure 8C and 9C). However, to demonstrate the effect of the implicit process switching the error-correction calculation to be consistent with the mirror reversal, we flipped the sign of the implicit error term in the Mirror-Task-Mirror-Correction Model (Figure 8D and 9D).

When the mirror task is simulated with error calculated as appropriate for a rotation [*aim – cursor*], the result is a response pattern much like that of the data. In contrast, an error calculation that is appropriate for the mirror [–(*aim – cursor*)], produces results in a radically different pattern from the data (compare Figure 8.A to 8.C-D and Figure 9.A to 9.C-D). The modeling results demonstrate that the implicit process under a mirror reversal first operates as if the perturbation was rotational in nature, then suppresses this adaptation.

In the washout phase, at the end of day five, subjects were instructed to aim directly at the target without using any strategy to measure the size of the aftereffect. Similar to Experiment One, we observed a very limited range of aftereffects compared with the implicit process calculated throughout training. After correcting for baseline bias, aftereffects hover slightly, yet significantly, below zero (average corrected aftereffect = −3.06 ± 3.2°, t-value =-3.32, p = .007, Figure 1). As in Experiment One, subjects were instructed to aim directly at the target without using any strategy in washout trials and no feedback was given. In addition to exhibiting the high within-subject variance seen in Experiment One, the washout phase was confounded by the discrepancy between aiming to hit a target in training versus aiming to hit that same target in washout. Adaptation appears to be centered around the aim location, not the target location (Day et al 2016; McDougle and Taylor, 2019; Schween et al 2019). In our task, a correct re-aiming location in training is approximately the same location as another target on the opposite side of the mirror axis.

The new aiming location in washout (i.e. directly at the target) is at the same location as that for the opposite target in training. Adaptation that resulted in positive aftereffects (relative to the adaptation expected in a visuomotor rotation task) during training would have resulted in negative aftereffects in washout. This suggests that we should read the average aftereffect as being in the direction to correct for a rotational error, not a mirror reversal.

#### Comparison of the Results of Experiments One and Two

Although Experiments One and Two were conducted sequentially, and therefore should not be statistically compared, we took care to keep both experiments as similar as possible to allow qualitative comparison.

We saw a steadily declining average implicit recalibration for subjects training to a mirror reversal in Experiment Two. This is in contrast to the flat average of the implicit process function seen for the duration of training to a visuomotor rotation. A visual comparison of these time courses shows starkly different behavior between the two groups (Figure 13A). While subjects completing the mirror reversal task did not, on average, reverse their implicit process sufficiently for it to work with their learning goal, they did suppress adaptation when compared to the subjects learning a rotation. This suppression is evident from the first day of training, where we see a slowing of initial adaptation rate. The source of this suppression remains an open question (see discussion).

**Figure 13.**
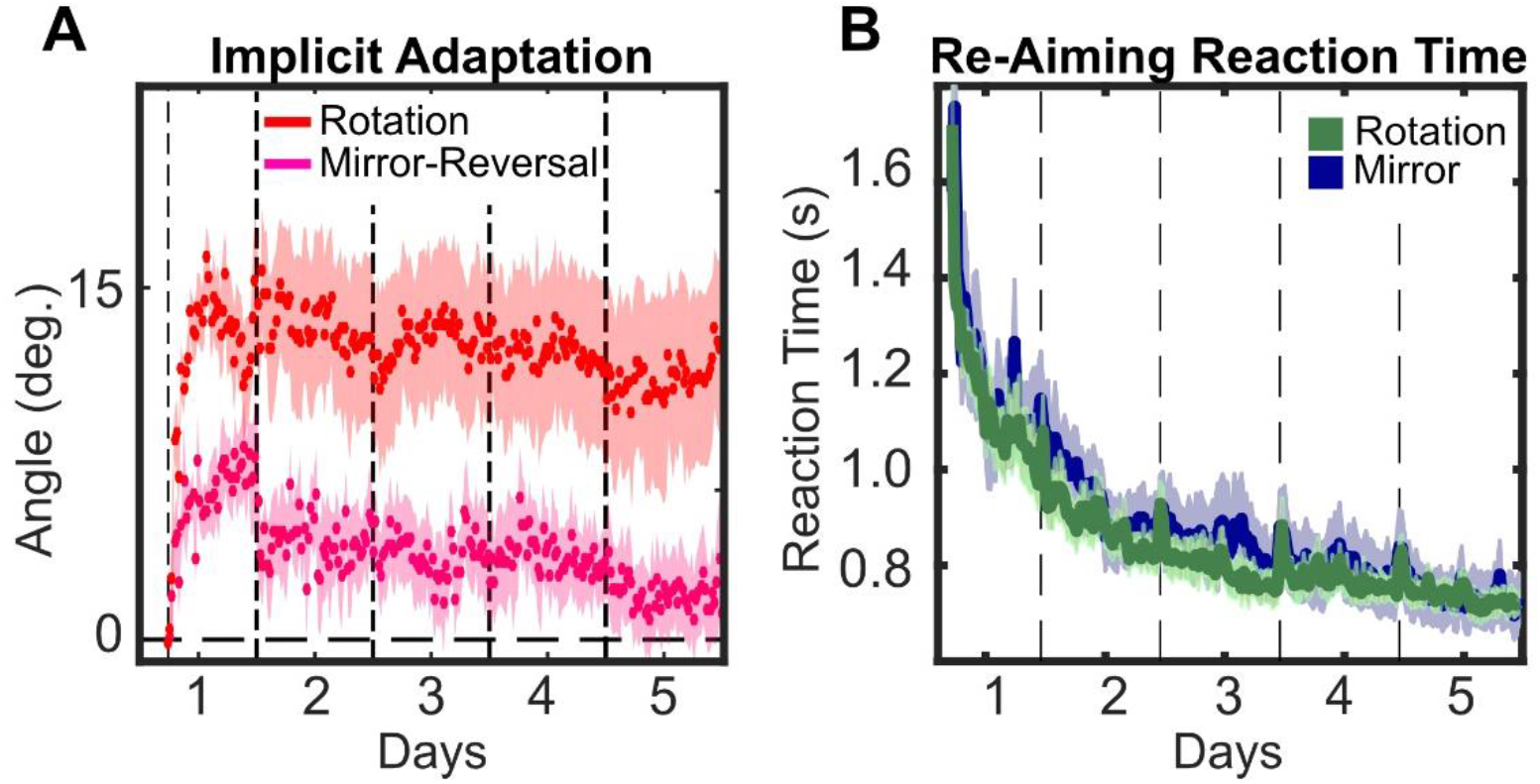
Comparison between subjects in Experiment One (Rotation) and Experiment Two (Mirror Reversal). The horizontal dashed lines represent first the onset of the perturbation, then day break. The shaded areas represent the standard error of the mean. A) Time course of calculated the implicit process. B) Reaction time across training calculated as interval from target onset to re-aim registered. Because re-aiming was introduced right before perturbation onset, this time-course starts when the perturbation was initiated.

We find that re-aiming reaction times, the time between target appearance and the touchscreen registering a re-aim tap, gradually decrease to approximately half over the course of five days of training (1.4 ± 0.2 s in first five bins of rotation training to 0.7 ± 0.07 s in last five bins of rotation training, Figure 13B). With no difference between the groups.

## Discussion

The idea that adaptation of the motor system to altered feedback is largely the result of implicit process can be dated as far back as Stratton (1896). Our everyday experiences reflect this view. For example, compensating for the lateral and vertical shift of a computer mouse or a new pair of eyeglasses do not feel like the result of explicit processes. However, previous studies that have sought to isolate implicit recalibration in visuomotor adaptation tasks have found that it appears to be limited (Bond and Taylor 2015; Morehead et al., 2017; Kim et al., 2018; Krakauer et al., 2019). Here, we sought to determine if the limited capacity of implicit recalibration could be overcome with extended training. In two experiments, we find that implicit recalibration to both a visuomotor rotation and mirror reversal remains limited, for the majority of participants, after five days of training.

### Limitation of Implicit Recalibration and Measurement Reliability

In these experiments, we measured the implicit process by subtraction from explicitly indicated re-aiming and as an aftereffect (Taylor and Ivry, 2014; Bond and Taylor, 2017),. Regardless of the metric used to measure the implicit process, we found that implicit processes do not fully account for learning in a rotation task. The implicit process in the mirror reversal task initially was counterproductive to performance before becoming suppressed with increased training. These findings may have been expected given the recent evidence that the implicit process falls well short of full learning in visuomotor adaptation tasks (Morehead and Smith, 2017; Morehead, Taylor, Parvin, and Ivry, 2017; Kim et al., 2018; Kim, Parvin, and Ivry, 2019) and in the wrong direction in the mirror reversal task (Gritsenko and Kalaska 2009; Lilicrap et al 2013; Telgen 2014; Kasuga 2015; Hadjiosif et al., 2021). Nonetheless, we thought that this limitation of implicit process may be overcome with more training, an idea that failed to be supported by our five-day experiments.

While group-level data does not provide evidence that the implicit process is sufficient for complete learning in a visuomotor rotation task, certain individuals do show complete adaptation under the subtraction measure. Enormous variability, spanning the entire range from zero to full adaptation through implicit processes, was seen across individuals. This suggests that the traditional method of averaging over participants is simplifying, and perhaps obscuring, experimental results (Gallistel 2004). It remains an open question as to why there is such a degree of inter-subject variability; however, this degree of variability has been observed in several studies where for even small rotations, on the order of 3-10°, they found implicit recalibration ranging from nearly zero to almost 50° (Stark-Inbar et al., 2017; Morehead et al 2107; Kim et al., 2018; Kim et al., 2019). Tsay and colleagues (2021) have found a modest yet significant correlation between the degree of proprioceptive variability and realignment with the degree of implicit recalibration. This finding is consistent with Bayesian sensorimotor integration framework where adaptation is a function of the weighting between of proprioception and vision based on their relative degrees of noise. Participants with more noise in their proprioceptive system may be more strongly driven by visual feedback, resulting in greater implicit recalibration for visuomotor perturbation.

An additional possibility for the limited degree of implicit recalibration observed here, as well as the variability between participants, may be the result of explicit strategy use. While several studies suggested a considerable degree of independence between implicit and explicit processes (Mazzoni and Krakauer 2006; Taylor and Ivry 2011; Taylor et al., 2014), this has recently been challenged. Participants who reported their explicit aiming direction throughout training, showed less implicit recalibration than participants who were only probed at the end of training (Maresch and Donchin 2020). By asking participants to report their aiming location on each trial, learning may be shifted more toward explicit strategies. It should be noted that in our original study of the aim report method, we found very little difference in the size of aftereffects between groups that reported on each trial versus never reported (Taylor et al., 2014). This was, in part, our reasoning for not including a no-report control group in conjunction with the difficulty of recruiting participants for a five-day study where the primary dependent measure (i.e., aftereffects) occurs only once at the end of the study. Clearly, more work is needed to flesh out this psychological equivalent of the measurement problem.

Finally, if implicit recalibration and explicit aiming operate on the same target error signal, then learning by one system would compete with the other (Albert et al., 2021). Indeed, there appears to be a reliable negative correlation between learning by explicit re-aiming and implicit recalibration (Albert et al., 2021); however, this is difficult to interpret since both processes are linked to cursor feedback and if implicit recalibration plateaued, then more explicit re-aiming would be required to restore performance. What’s more, the initial findings by Mazzoni and Krakauer (2006) strongly suggested that implicit recalibration operated on a sensory-prediction error and not target error, but this has recently been challenged (Ranjan and Smith 2020; Albert et al., 2021).

### Implicit recalibration in the mirror reversal task

The mirror reversal results present an exciting future opportunity to determine the error signal that initiates suppression of the implicit process in the mirror reversal task. Large variability in individual behavior provides a foothold to understanding suppression of implicit processes. Here we consider a few possible explanations that could be a target for future research.

The mirror reversal, unlike a rotation task, has an error-signal with a constantly switching sign. Previous work has shown that inconsistency in error signals reduces overall learning (Castro et al., 2014) and the implicit process in particular (Hutter and Taylor, 2018), which is consistent with the motor system storing a history of errors (Herzfeld et al., 2014). Here, we would expect participants that experienced more error switching, through their unique history of missing the target, to suppress implicit adaptation more quickly and fully (Albert et al., 2020).

Second, the difference between rotations and mirror-reversals could be thought of as a reparameterization, as previously considered in rotations, compared with learning a completely new structure, suggested for the mirror reversal (Braun, Aertsen, and Wolpert, 2009). The reparameterization of the transformation matrix, as seen in adaptation to a rotation, might be the fundamental function of the implicit process. Indeed, the implicit process may be exceptional at matrix reparameterization. In contrast, learning a mirror reversal requires learning the full structure of the task, including the relationship between parameters in the transformation matrix. It appears that the implicit process tries to reparametrize the transformation matrix as if it were a rotation even if doing so is inappropriate for the task. Only after days of training does the implicit process appear to stop reparametrizing inappropriately. It could be that once a new structure is specified, the structure of the mirror reversal, the implicit process becomes appropriate for this new structure. The speed of learning this structure would dictate how quickly implicit processes began reparametrizing appropriately.

A slightly lower-level of description of this problem, one rooted in control engineering, was put forth in a recent study by Hadjiosif and colleagues (2021), which compared adaptation under a visuomotor rotation and a mirror reversal as a way to distinguish between updating forward model updating and updating a control policy. They too find that the implicit process in a mirror reversal was directed in the wrong direction to counter a mirror reversal, which is inconsistent with the updating a forward model. Instead, implicit learning of a mirror reversal may require direct updating of a control policy. Learning a new control policy would again have the outcome of changing the structure of the transformation matrix.

These explanations are highly speculative, and additional research is necessary to pin down the reason for implicit processes being counterproductive in a mirror reversal task, and the subsequent suppression of implicit recalibration.

### Proceduralization of Explicit Re-aiming

Although the implicit process is insufficient for learning, we submit that it is highly unlikely that subjects are performing a time-intensive, computationally-demanding, strategizing process at the end of five days of training. A more likely explanation is that the re-aiming strategy has become partially proceduralized, also referred to as caching or habitualization (Huberdeau et al., 2019). We would expect to find evidence for proceduralization in participants’ reaction or preparation times (Logan, 1980; Cohen, Dunbar, and McClelland, 1990). Increased reaction time increases are commonly observed at the onset of visuomotor perturbations and gently decline with training (Saijo and Gomi, 2010; Fernandez-Ruiz, Wong, et al., 2011; McDougle and Taylor, 2019). If preparation time is limited, performance is significantly hindered in visuomotor rotation tasks as participants may not have sufficient time to re-aim their movements to counter the rotation (Fernandez-Ruiz et al., 2011; Haith et al., 2015). However, one study found that performance under constrained preparation time can be restored following two days of training, potentially reflecting a proceduralization of the re-aiming strategy (Huberdeau, Krakauer, and Haith 2019).

Here, we find that re-aiming reaction times in both experiments gradually decrease to approximately half over the course of five days of training (1.4 ± 0.2 s in first five bins of rotation training to 0.7 ± 0.07 s in last five bins of rotation training, Figure 13B). We would like to interpret this result as indicative of proceduralization of the re-aiming strategy in both tasks. Note: the absolute value of reaction time during training is expected to differ from previous reports, as our measure of reaction time necessarily includes the time taken to report the re-aiming location with the non-dominate hand (e.g. Fernandez-Ruiz, Wong, et al. 2011; Haith, Huberdeau, and Krakauer 2015). Nonetheless, the fact that we observe a significant drop in reaction times over the course of training, and given the traditional view of interpreting reaction times as reflective of computational demands, we suspect that this is an indication that proceduralization has occurred.

The question turns to what might underlie proceduralization of explicit re-aiming. One possibility is that the process is as simple as forming a stimulus-response mapping, allowing subjects to automatically perform an action that was previously conducted through computation (Daw, Niv, and Dayan, 2005). Additionally, stimulus-response mappings may result in a cached response or habit, which could simulate an implicit process by being extremely fast and relatively robust to added cognitive loads (Daw, Niv, and Dayan, 2005; Dayan, 2009; Haith and Krakauer, 2013). McDougle and Taylor (2019) found evidence that initial rotation re-aiming strategies eventually give-way to a theorized look-up table for each target location (a stimulus-response map), echoing Logan’s dual process theory of skill as reflecting a shift from algorithmic-based to retrieval-based operations (Logan 1988).

### Summary

In these two experiments, we found that, on average, the implicit process does not account for the majority of adaptive learning, even after many days of consolidation. However, we also saw large variation between individual subjects. This variation questions the validity of averaging over subjects to make claims about implicit processes, and raises questions as to why one measure captures this variability while the other does not. Finally, we found evidence in the mirror reversal task that the implicit process is slowly suppressed to compensate for alternate task demands.

## Acknowledgements

We thank Peter Butcher for providing extensive coding assistance in preparing the main experimental task.

